# PROTEOME-SCALE RECOMBINANT STANDARDS AND A ROBUST HIGH-SPEED SEARCH ENGINE TO ADVANCE CROSS-LINKING MS-BASED INTERACTOMICS

**DOI:** 10.1101/2023.11.30.569448

**Authors:** Milan Avila Clasen, Max Ruwolt, Louise U. Kurt, Fabio C Gozzo, Shuai Wang, Tao Chen, Paulo C Carvalho, Diogo Borges Lima, Fan Liu

## Abstract

Advancing data analysis tools for proteome-wide cross-linking mass spectrometry (XL-MS) requires ground-truth standards that mimic biological complexity. Here, we develop well-controlled XL-MS standards comprising hundreds of recombinant proteins that are systematically mixed for cross-linking. We use one standard dataset to guide the development of Scout, a search engine for XL-MS with MS-cleavable cross-linkers. Using other, independent datasets from our standards as well as published datasets, we benchmark the performance of Scout and existing XL-MS software. This demonstrates that Scout offers the best combination of speed, sensitivity, and false-discovery rate control. These results illustrate how our large recombinant standards can support the development of XL-MS analysis tools and evaluation of XL-MS results.

## Introduction

Cross-linking mass spectrometry (XL-MS) is a powerful technique to analyze protein structures and interactions by providing residue-to-residue connections at low-nanometer resolution^1^. Over the last decade, the scope of XL-MS has expanded from purified proteins/complexes to (sub)proteomes. Two main driving forces of this development were the introduction of MS-cleavable cross-linkers and the development of advanced cross-link search engines^2-5^. Cross-linkers that are cleaved during MS analysis produce linear peptides with signature fragmentation patterns that allow discerning cross-link mass spectra from others, deriving the mass of the linked peptides, and – through further fragmentation – sequence the individual peptides. Consequently, the search space increases linearly instead of quadratically with the number of protein sequences in the database, which particularly benefits full proteome analyses and is leveraged by multiple search engines.

Benchmarking XL-MS search output is traditionally achieved by taking three-dimensional protein structures as the ground truth and testing how well they match the identified crosslinks^6^. However, this approach was shown to substantially underestimate the false-discovery rate in proteome-wide XL-MS studies^7^. More recently, a synthetic XL-MS peptide standard has been developed to serve as a ground truth for benchmarking XL-MS search engines^8^, but this standard only comprises 141 peptides from 38 proteins and thus is too small to provide insights into software performance in proteome-wide XL-MS experiments. Establishing software tools for proteome-wide XL-MS has primarily relied on *ad hoc* produced biological samples (e.g., fractionated *E*.*coli* lysate^9, 10^ and spiked-in ^15^N metabolically labeled datasets^11^) or transfer learning of models originally trained on linear peptides^12^, but the lack of an analytical XL-MS standard that can mimic complex biological samples has hindered their unbiased validation and benchmarking. An additional challenge for XL-MS software benchmarking arises from the fact that, depending on the specific application of XL-MS, identifications need to be reported on three different levels referred to as cross-link spectrum matches (CSMs), unique residue pairs (ResPairs), and protein-protein interactions (PPIs). An ideal XL-MS standard should provide a reliable ground-truth at all these levels.

Here, we provide an analytical XL-MS standard that is an order of magnitude more complex than existing ones. We produced hundreds of human proteins or protein fragments in *E. coli* and cross-linked them according to a pre-defined mixing scheme, resulting in fully controlled XL-MS datasets. One dataset guided the development of Scout, an FDR-controlled cross-link search engine that relies on artificial neural networks (ANN) and is optimized for XL-MS with MS-cleavable cross-linkers. The other, independent standard datasets were used as a ground-truth to benchmark Scout and several state-of-the-art search engines (xiSEARCH/xiFDR^13, 14^, MaxLynx^15^, MSAnnika^4^, XlinkX in Proteome Discoverer v2.5^16^ (XlinkX PD), MeroX^17^) at CSM, ResPair and PPI level in differently sized search spaces. The results show that Scout is unique in its ability to unite high sensitivity (high number of true identifications), specificity (accurate FDR) and speed (low processing time). This holds true across all levels of cross-link identification and for all database sizes we have tested. Scout also performs well on previously published datasets for FDR estimation^8, 10^, yielding more identifications than the best-performing XL-MS search engines in these previous publications. These data demonstrate the power of Scout as well as the potential of our large-scale XL-MS standards to support the development of machine learning-based methods and other algorithmic solutions for analyzing XL-MS data from complex samples.

## Results

### Constructing an XL-MS standard mimicking complex biological samples

Our XL-MS standard is based on human proteins or protein fragments produced in *E. coli* (see Supplementary Data 1 for a full list and SDS-PAGE quality control). We divided them into 32 interaction groups with 8 proteins each and cross-linked each group separately with DSSO under conditions to maximize cross-link formation and consequently the number of detectable CSMs (see Methods, Figure 1). Proteins from the same interaction group were mixed pairwise in all possible combinations, cross-linked, and combined into a pooled sample for digestion, strong cation exchange fractionation and LC/MS analysis. As a result, each protein will be inter-linked to 7 pre-defined interactors, allowing up to 896 unique PPIs. Any inter-links between proteins within the same interaction group that a cross-link search engine may identify are considered true-positive hits; any identified inter-links between proteins from different groups must be false-positives. We also consider any intra-links within proteins as true-positives, since they are much less prone to arise from random matching, even with increasing search space, and are therefore unlikely to be false-positives^2, 10^.

**Figure 1.**
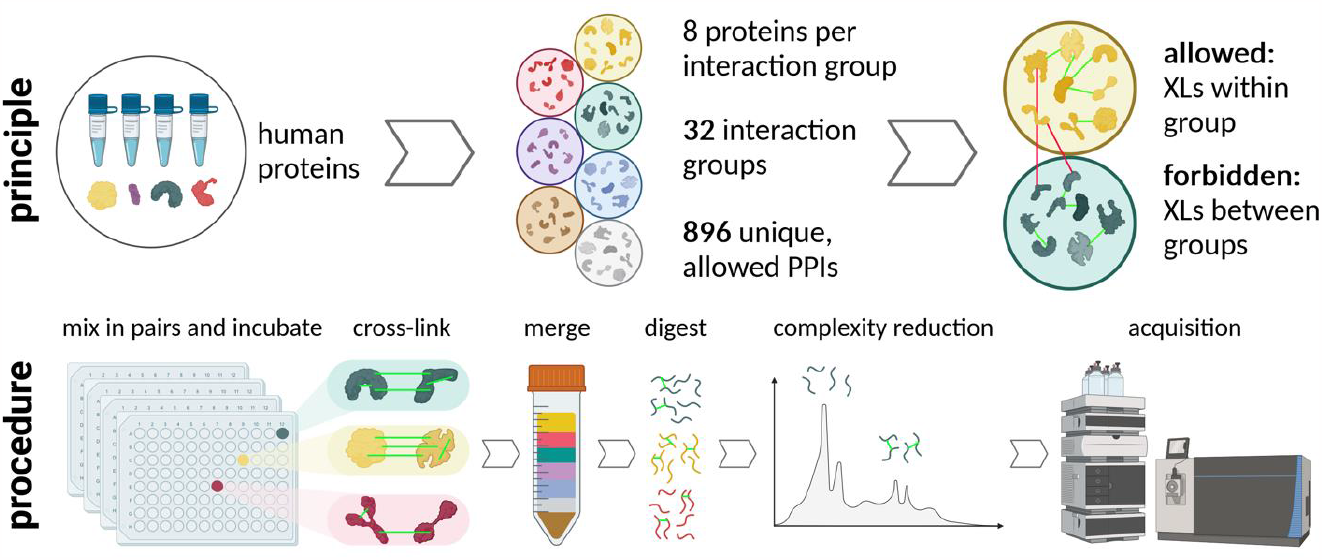
Schematic workflow for the construction of the XL-MS standard. Proteins were allocated into 32 interaction groups with 8 proteins each. Within the interaction groups, proteins were cross-linked pairwise in all possible combinations, resulting in 28 PPIs per interaction group and 896 PPIs in total. All cross-linked samples were merged prior to digestion.

### Using the XL-MS standard to develop an artificial neural network-based cross-link search engine

Our well-controlled XL-MS standard yields datasets that can aid the development of algorithms for XL-MS data analysis. As a first example, we introduce Scout, a cross-link search engine for identifying peptides cross-linked with cleavable reagents. Scout is an intuitive, userinterface-controlled software that relies on artificial neural networks (ANNs) to generate discriminant functions to score and rank identifications using several quality metrics optimized at the CSM, ResPair and PPI levels. Scout enables multi-tier FDR filtering at all levels. Scout’s workflow is shown in Figure 2 and described in detail in Supplementary Notes 1-2.

We used one standard XL-MS dataset (based on 8 interaction groups) to guide the development of the ANNs (Supplementary Notes 3-4). This subset of our standard comprises 124,794 identified target CSMs and 149,638 decoy CSMs. This CSM-level depth is similar to a recent XL-MS study in intact human cells^18^, increasing our confidence that this dataset can mimic a proteome-wide XL-MS experiment.

**Figure 2.**
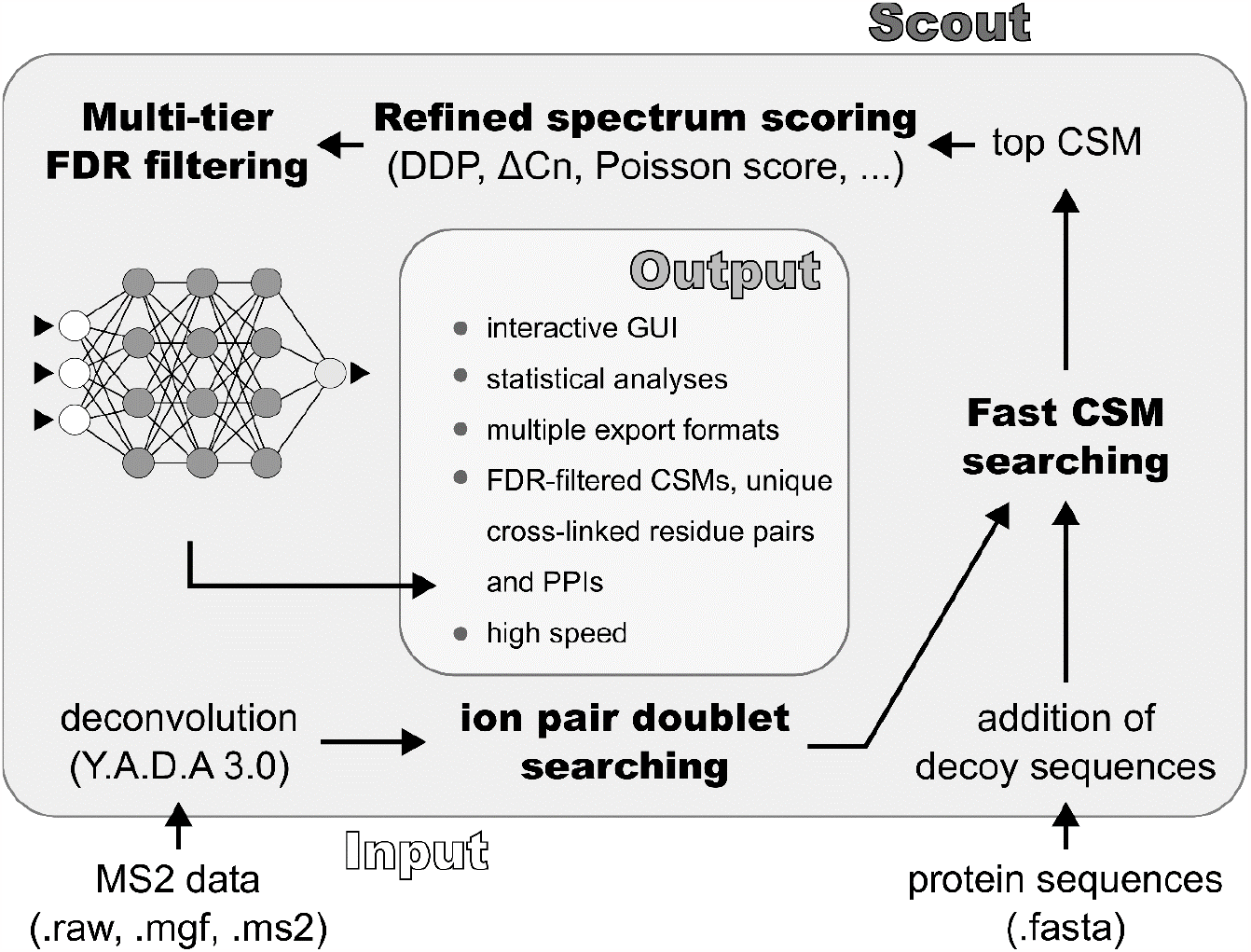
Schematic representation of the cross-link identification workflow employed by Scout. Scout requires mass spectrometry raw data (MS2 spectra) and a protein sequence database as input. Cross-links are identified in two search steps – ion pair doublet searching and fast cross-link spectrum match (CSM) searching – which are both described in Supplementary Note 1. The shortlisted peptide pair candidates for each MS2 spectrum are then subjected to refined spectrum scoring based on a set of sensitive quality metrics described in Supplementary Note 2. Finally, the results are filtered according to a user-defined FDR using a machine learning-based discriminant function at each tier of identification - CSMs, ResPairs and PPIs (see Supplementary Note 3). The final output is presented through a graphical user interface, providing a user-friendly display of the identified cross-linked peptides and their associated metrics.

### Benchmarking Scout on independent XL-MS standard datasets

In addition to aiding software tool development, the datasets derived from our standard can serve as a ground-truth: We know for each detected cross-link whether it is allowed (inter-links and intra-links within one interaction group) or not allowed (between-group inter-links, or intralinks of proteins not present in the respective group) according to our mixing scheme (Figure 1). This information enables us to calculate an empirical FDR similar to a target-decoy FDR^13^, which offers the opportunity for an unbiased benchmarking of XL-MS search engines.

We compared Scout with the following widely adopted XL-MS search engines that are compatible with MS-cleavable cross-linkers: MaxLynx, MSAnnika, XlinkX PD, MeroX, and xiSEARCH with the xiFDR module. We benchmarked these tools with those standard datasets that were not used during Scout development. To avoid that a protein was assigned to the wrong interaction group because of non-unique peptides within the dataset, we allowed more than one possible group per peptide if the peptide was shared between homologous proteins.

We varied the search space size, using a small 540-protein database a large 4,000-protein database, both comprising the proteins present in our standard and randomly selected human entrapment proteins from SwissProt with < 85% sequence identity to our recombinant proteins (see Supplementary Table 1). We selected the same search parameters for each software wherever possible, and set a 1% FDR cutoff within each software (referred to as naïve FDR) with separate FDR filtering for inter- and intra-links. If a software tool did not provide naïve FDRs on all identification levels (CSM, ResPair, PPI), we took the naïve FDR-filtered identifications from the next lowest level and aggregated them to the higher level without any post-processing filters and score cutoffs (referred to as “aggregated results”). A full description of the search settings is provided in Supplementary Table 2. For clearer visualization, results are presented separately for inter-links (Figures 3, 4) and intra-links (Supplementary Figure 1). XlinkX PD and xiSEARCH/xiFDR had to be analyzed separately as explained below.

For intra-links, all search engines properly controlled the FDR below 1% on CSM and ResPair level, except for MeroX (> 20%). Since intra-links are used for investigating protein structure rather than PPIs, the most relevant information comes from the ResPair level, for which Scout shows the best overall performance (Supplementary Figure 1).

For inter-links, Scout outperformed MaxLynx, MSAnnika, and MeroX on CSM and ResPair levels, yielding smallest empirical FDR and highest true-positive identification numbers irrespective of the database size (Figure 3). At PPI level, MSAnnika reports more true-positive PPIs than Scout, but at a substantially higher empirical FDR (12-15%, Figure 3 – right panels), which is expected because MSAnnika does not include a dedicated FDR control at PPI level. This emphasizes the importance of controlling FDR on all identification levels^13^.

**Figure 3.**
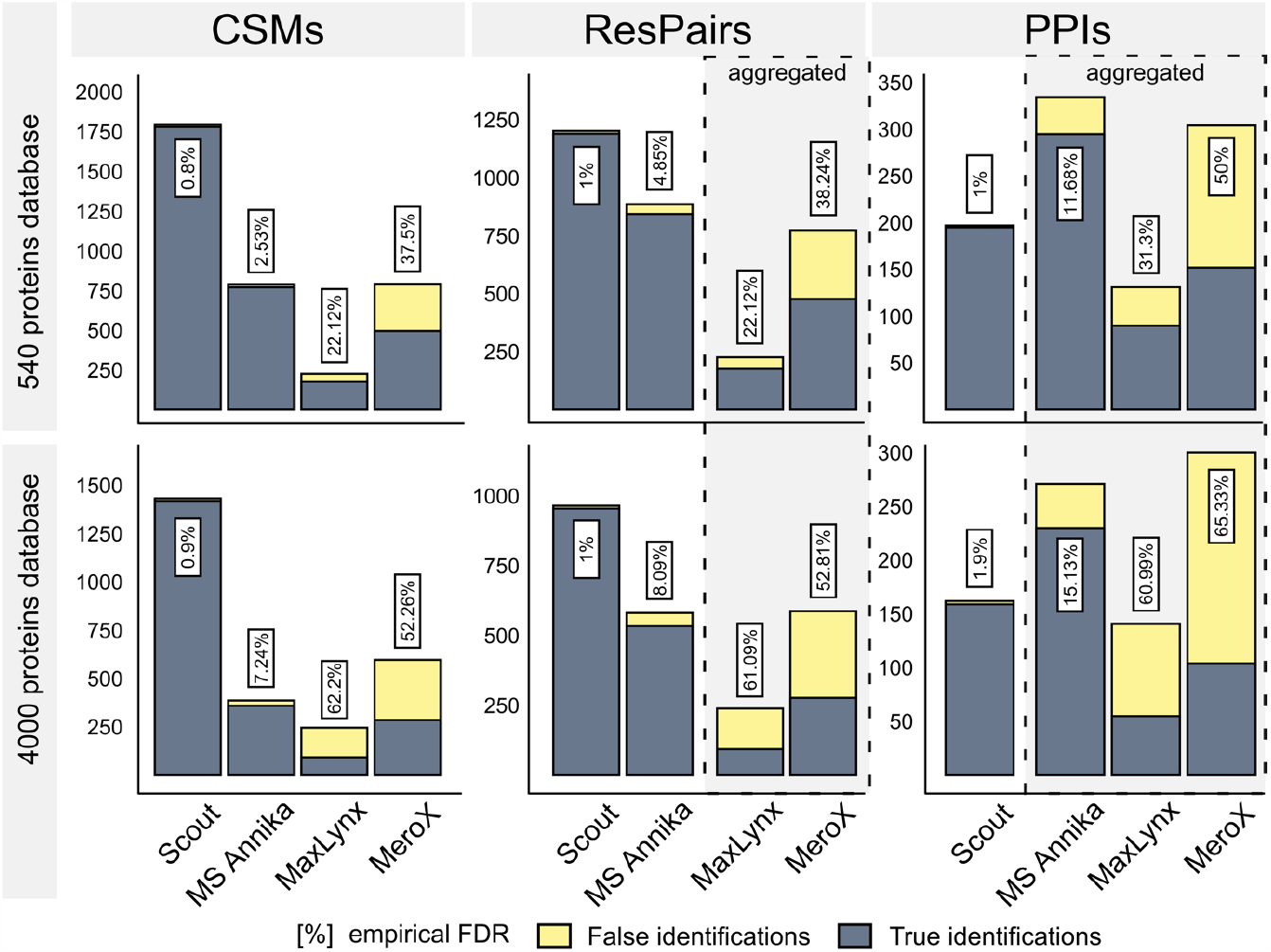
Benchmarking Scout against other XL-MS search engines for inter-link identifications. Number of identified inter-protein CSMs, ResPairs and PPIs and the empirically determined FDR at a naïve 1% FDR cutoff using a 540-protein or 4,000-protein database and identical search parameters. Framed bars mark aggregated results, i.e. cases when CSMs were aggregated to unique ResPairs or unique ResPairs to unique PPIs because search engines do not control FDR at these levels (MeroX and MaxLynx do not report FDR-controlled ResPairs; MeroX, MaxLynx, MSAnnika do not report FDR-controlled PPIs). Blue bars show true-positive identifications, yellow bars show false-positive identifications, violating the mixing scheme of our XL-MS standard. The empirical FDR is shown in boxes above the bars.

The inter-link comparison of Scout and XlinkX PD was done separately, because the XlinkX PD output is substantially affected by post-processing settings that are not part of our standard search parameters. In particular, the FDR in XlinkX PD is substantially affected by heuristic score cutoffs, which were shown to strongly depend on dataset and search parameters^6^. A recent study suggested a minimum XlinkX score of 60^19^, whereas the XlinkX PD user guide proposes to set dynamic score cutoff based on the score of the best CSM-level decoy hits in each analysis (see Methods). We tested both of these options (Figure 4A-B) as well as score cutoffs that would filter the results to 1% empirical FDR (Figure 4C). With regard to identification numbers, Scout outperformed XlinkX PD on CSMs and ResPairs, whereas XlinkX PD identified more true-positive PPIs in most settings. However, with regard to identification confidence, XlinkX PD strongly depends on manually setting suitable score cutoffs during post-processing while Scout is able to maintain an empirical FDR <2% on all identification levels (Figure 3). Furthermore, XlinkX PD identifications are more susceptible to entrapment. At 1% empirical FDR, XlinkX PD reports a higher number of PPIs than Scout when searching against the 540-protein database, whereas Scout outperforms XlinkX at the 4000-protein database search (Figure 4C). These results suggest that Scout is more robust and suitable for identifying PPIs against large sequence databases.

The comparisons of Scout to xiSEARCH/xiFDR were also performed sepearately because searches against the small 540-protein database with xiSEARCH/xiFDR did not run to completion after several weeks when using our standardized settings and server equipment. This is in line with previous reports that proteome-wide applications of xiSEARCH/xiFDR on in-house computers are highly restricted^8^. To still compare Scout to xiSEARCH/xiFDR, a subset of the bechmarking dataset was run on a computer cluster using more restrictive developer-recommended parameters (see Methods) and the 540-protein database. Scout (run with the same parameters as in Figure 3) reported more correct identifications on all levels when filtering data with a naïve 1% or 5% FDR cutoff (Figure 4D, Supplementary Figure 2A).

We also compared the data processing times of all tested software on a server equipped with 512GB RAM and powered by dual Intel(R) Xeon(R) Gold 6136 CPUs operating at 3.00GHz. Scout was substantially faster than the other tools in small and large database searches, and showed the smallest speed decline with increasing database size (Figure 4E). To compare the processing time of xiSEARCH and Scout using the same computational setup, we analyzed four RAW files of the benchmarking dataset with Scout’s and xiSEARCH’s default parameters and the 540-protein database (Supplementary Figure 2B). Scout processed the data >200-times faster than xiSEARCH. Importantly, while this high-capacity server was needed to meet the RAM demands of some of the tested tools, Scout operates efficiently with a small memory footprint and is well-suited for desktop PCs with as little as 16GB RAM (see also Figure 5B). Thus, Scout offers the best combination of sensitivity, specificity and speed on all three levels of XL-MS identifications irrespective of the search space.

**Figure 4.**
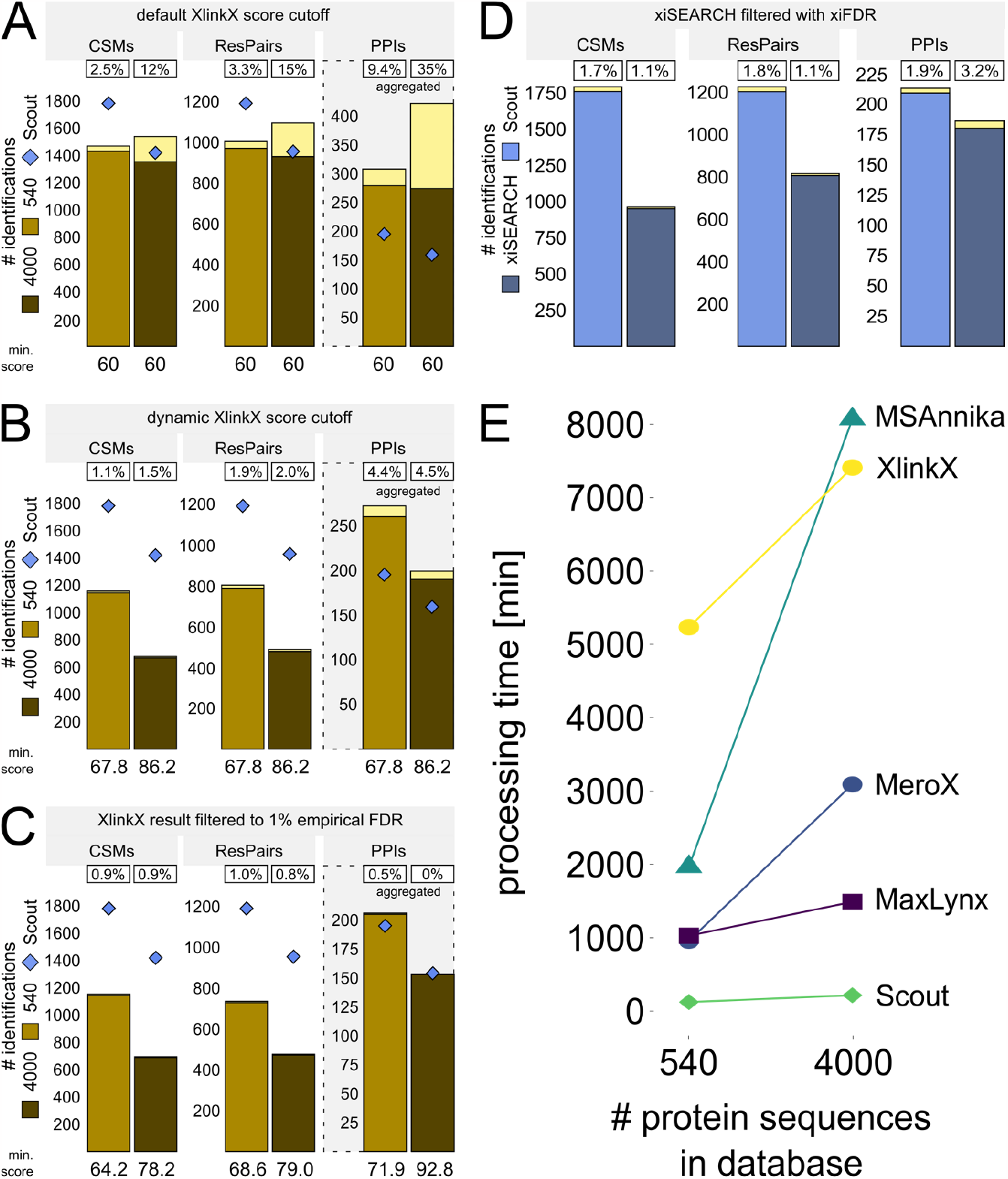
Benchmarking of XlinX PD, xiSEARCH/xiFDR, and software processing times. (A-C) Inter-protein CSMs, ResPairs and PPI identifications when comparing Scout and XlinkX PD. Truepositive identifications by XlinkX PD are shown in light-brown (540-protein database) and dark-brown (4000-protein database). The Scout numbers (blue diamonds) are the same as in Figure 3. In addition to using our standard search parameters, XlinkX PD identifications were post-processed using (A) previously optimized score cutoffs (‘default’) ^19^, (B) score cutoffs derived from the highest-scoring CSM-level decoy in every analysis (‘dynamic’), and (C) score cutoffs set to filter XlinkX PD results to 1% empirical FDR. The XlinkX score cutoffs are displayed below the bars. (D) Inter-protein CSMs, ResPairs and PPI identifications when comparing Scout and xiSEARCH. For Scout, results were filtered at 1% naïve FDR on all levels. For xiSEARCH/xiFDR, following the developer’s recommendation, a 1% naïve FDR was applied only on PPI level using boost between proteins (xiFDR) and reported are the resulting PPIs together with their corresponding CSMs and ResPairs. Scout and xiSEARCH were run using their default parameters, respectively, with KSTY as the possible reaction sites for the cross-linking reagent. In panels A-D, the framed percentage numbers indicate the final empirical FDR and yellow bars show false-positive identifications, violating the mixing scheme of our XL-MS standard. (E) Processing time using different search engines on the benchmarking dataset with a 1% naïve FDR cutoff.

Next, we tested how increasing the search space and adding entrapment sequences impacts Scout’s overall performance when using standard parameters and a 1% naïve all-level FDR cutoff. Scout maintains a low empirical FDR on all levels at most tested database sizes (Figure 5A). The number of identified inter-protein CSMs, ResPairs and PPIs decreases as expected. For example, moving from an entrapment 4-times higher than the number of experimentally available proteins to a 160-times higher entrapment reduces PPI identifications by 23%. However, our subsequent analyses indicate that this decrease is less drastic than for other search engines (see Supplementary Table 3 and related discussion below). Meanwhile, the search of 55 RAW files with 540 protein entries takes only 2h on a desktop computer (Intel Core i7 2.90GHz, 16GB RAM). When searching the same RAW files against a 35-times larger database (20622 protein entries), Scout shows an acceptable 4.7-fold processing time increase to ∼9.5h, demonstrating that it operates efficiently even in large search spaces and is compatible with a standard desktop PC (Figure 5B).

**Figure 5.**
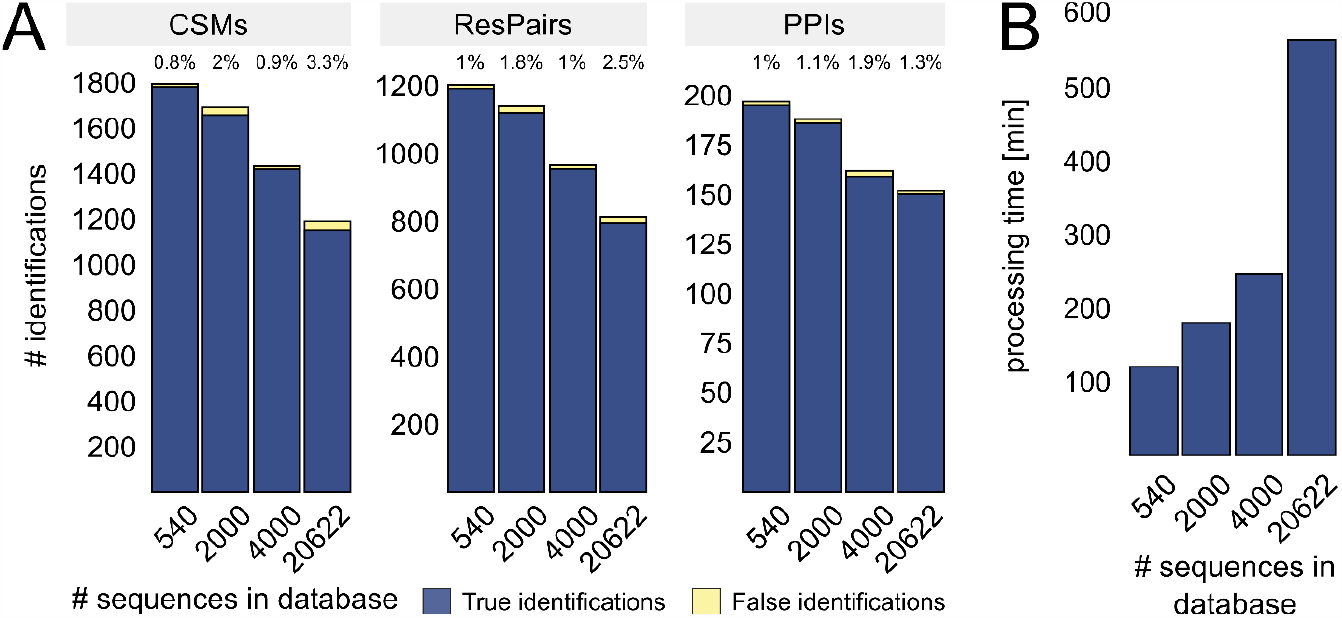
Search space-dependent performance of Scout. (A) Number of inter-protein CSMs, ResPairs and PPIs identified by Scout using standard parameters and all-level 1% naïve FDR cutoff on the benchmarking dataset with increasing database size. True positives shown in blue, false positives in yellow. Empirical FDR is indicated above the bars. (B) Processing time of Scout when searching the benchmarking data against increasingly large databases.

We further evaluated Scout’s performance on a published small-scale dataset. ^8^ Here, a synthetic peptide main library was cross-linked with DSSO according to a mixing scheme prior to mixing them with non-cross-linked tryptic HEK cell peptides at a 1:5 ratio. This allows FDR assessments at the CSM and ResPair levels^8^ with a realistic background signal arising from linear peptides as one would expect from complex interactomics experiments. We compared Scout to the best-performing software reported in the original publication (MSAnnika). Using the same search parameters as above and the database(s) provided in the original publication, we obtained highly similar ResPair-level results for both tools (Figure 6A). For the nonoverlapping identifications, however, Scout achieves a lower empirical FDR.

Again, Scout maintains its efficiency and speed in a scenario with higher entrapment: Increasing the search space shows that larger database sizes expectedly reduce the number of ResPair identifications but do not substantially impact the FDR or processing time (Figure 6B), confirming the results obtained with our own XL-MS standard (Figure 5). Overall, Scout identifies more unique true positive ResPairs than all search engines reported in the original paper (Supplementary Table 3). ^8^ Our software maintains the lowest FDR and highest identification numbers for database searches against 671–20,334 protein entries.

Finally, as the dataset from Matzinger et al. ^8^ does not provide PPI-level information, we turned to the dataset from Lenz et al. ^10^, which aims to provide a PPI-level quasi-ground-truth based on the abundance of proteins in cross-linked SEC fractions of *E. coli* lysate. As this dataset was used to advance xiSEARCH/xiFDR, we compared Scout to the xiSEARCH result reported in the original publication. On all identification levels, both software tools show a low empirical FDR (following the definition from the original publication) but Scout provides more identifications (Figure 6C-D). Scout identifies 24% more PPIs, but the overlap of PPI identifications between both search engines is modest (Figure 6C).

**Figure 6.**
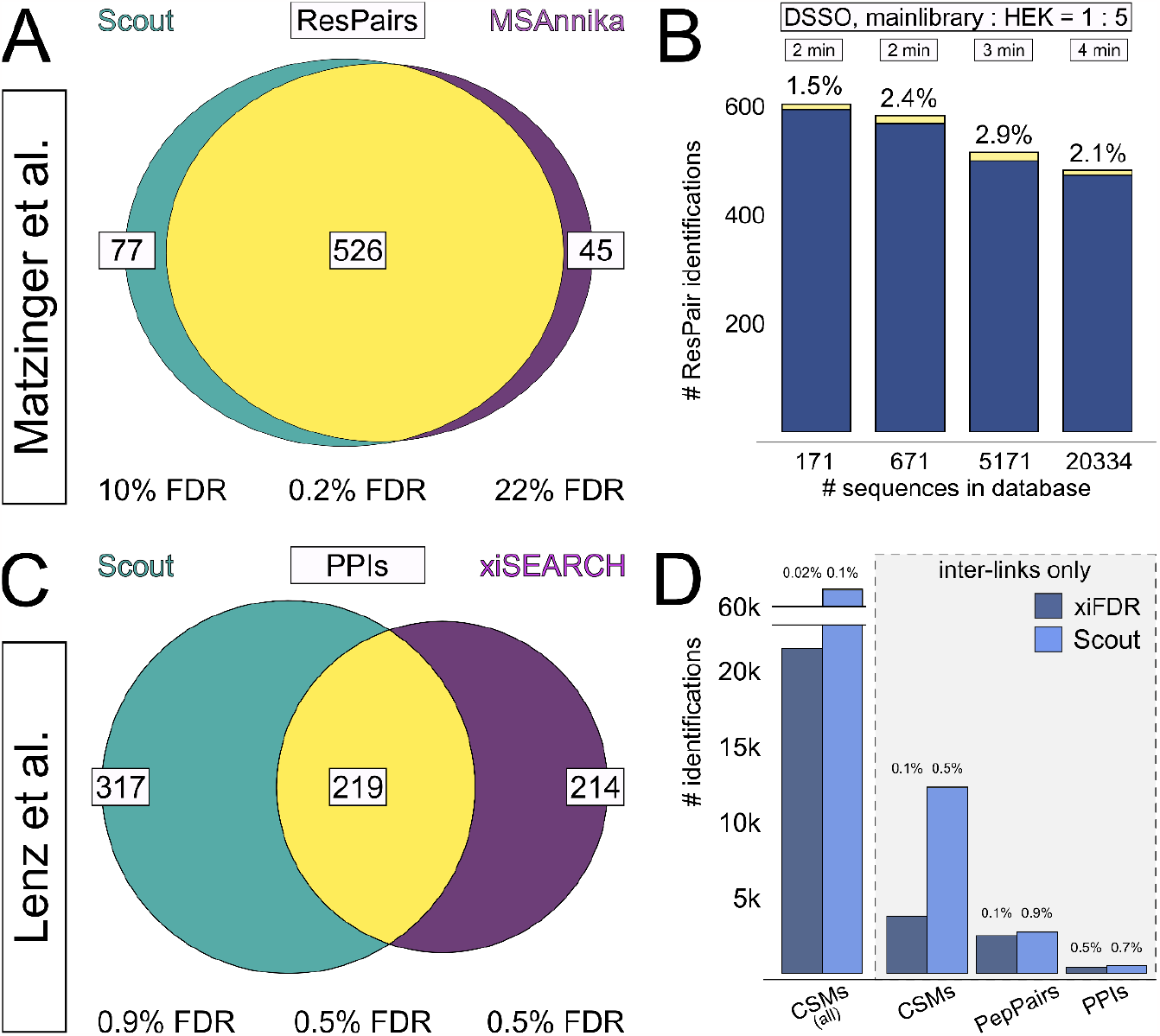
Performance of Scout on published XL-MS benchmarking datasets from Matzinger et al. using synthetic peptides^8^ (A, B) and Lenz et al. using fractionated *E. coli* lysate^10^ (C, D). (A) Overlap of ResPairs identified by Scout and MSAnnika and the true FDR of Scout-specific (left), shared (middle) and MSAnnika-specific (right) identifications using the DSSO main library spiked 1:5 into tryptic HEK peptides by Matzinger et al. using our standard parameters, which are similar to the ones reported in the original publication. (B) Scout’s true-positive (blue) and false-positive (yellow) ResPair-level identifications on increasingly large databases. The naïve FDR cutoff was set to 1%; empirical FDR and operating times are indicated above the bars. (C) Overlap of PPI-level identifications from Scout (left) and xiSEARCH (right) using the PPI benchmarking dataset by Lenz et al. and 1% separate naïve FDR cutoff on PPI level. Scout was operated with standard parameters and xiSEARCH identifications were retrieved from the original publication. Empirical FDR was determined using the procedure suggested in the original publication. ^10^ (D) Performance of Scout and xiSEARCH in identifying intra- and inter-protein CSMs, inter-protein CSMs only, inter-protein PepPairs and PPIs when setting an all-level naïve FDR cutoff of 1%. Scout was operated with standard parameters and xiSEARCH identifications were retrieved from the original publication. The empirical FDR was calculated as described by Lenz et al. and is indicated above the bars.

## Discussion

Advancing methods for proteome-wide investigations critically depends on the availability of thoroughly characterized analytical standards that provide a ground truth to validate new workflows and technologies. This is exemplified by the ProteomeTools project^1^, a synthetic peptide library that covers the entire human proteome and important post-translational modifications. ProteomeTools has become an invaluable ground-truth training dataset that has enabled the development of deep learning architectures to predict peptide tandem mass spectra, collision cross sections, and chromatographic retention times^20^. Related efforts for XL-MS have remained at relatively small scale, only covering a few dozen proteins^8^, which is too small to resemble proteomic complexity. Here, we mitigate this limitation by formulating a robust XL-MS standard that is an order of magnitude more complex. By systematic mixing and controlled cross-linking of our recombinant proteins, we generated a ground-truth of allowed and prohibited cross-link contacts that achieve the CSM-level scope of published proteome-wide XL-MS studies.

To demonstrate the utility of the resulting standard datasets, we used one of them to develop ANN modules for Scout, a new search engine for XL-MS studies with cleavable cross-linkers. Our standard dataset allowed us to fine-tune the sensitivity of Scout’s ANNs and avoid overfitting when training them on different identification levels. However, it should be noted that the number and mixing scheme of the proteins in our XL-MS standard does not resemble a naturally occurring PPI network. Hence, tuning PPI-level performance of XL-MS software tools can likely be improved further by designing larger standards with more complex mixing schemes that show greater similarity to PPI networks from different biological samples.

Nonetheless, our standard datasets are sufficient to calculate an empirical FDR at PPI level, which is a distinct advantage over the published small-scale XL-MS standard^8^. Leveraging this advantage, we benchmarked Scout’s performance against state-of-the-art XL-MS tools on CSM, ResPair, and PPI level. Importantly, this was done with datasets that had not been used during Scout development. These results – as well as performance comparisons on published benchmarking datasets^8, 10^ – demonstrate that Scout provides the best combination of accuracy, sensitivity, specificity and speed.

Overall, we believe that the fast and robust cross-link identifications enabled by Scout have the potential to directly advance proteome-wide XL-MS studies, while the large-scale XL-MS standard we provide can serve as a basis for developing a new generation of analysis tools for proteome-wide XL-MS.

## Methods

### Protein mixing and cross-linking

Ectopically expressed and purified human proteins were provided by Absea Biotechnology Ltd., Beijing (Supplementary Data 1). In brief, all proteins are His-tagged at the C-termini, packaged in PC3.1 expression plasmid, and produced in *E*.*coli*. Lyophilized proteins were dissolved in 20 mM HEPES pH 7.8 at concentrations varying from 0.1–1.5 mg/mL. The dissolved proteins were mixed in pairs of two proteins in all possible combinations within one interaction group. They were incubated for 20 min at 50 °C to induce interactions *in vitro*. 0.2– 1 mM DSSO cross-linking reagent was added to the groups and incubated at room temperature for 30 min. The cross-linking reaction was quenched with 20 mM Tris-HCl pH 8.0 for 30 min at room temperature. Subsequently, all groups were merged, and 8 M Urea was added to the solution. The protein mixture was reduced with 5 mM DTT for 1 h at 37 °C, alkylated in the dark with 40 mM CAA for 30 min at room temperature and digested with 1:200 (wt/wt) Lys-C for 4 h at 37 °C, shaking. After dilution with three volumes 20 mM HEPES pH 7.8, 1:100 (wt/wt) Trypsin was added and incubated over night at 37 °C. The digestion was stopped by adding 1% FA. Peptides were desalted with Sep-Pak C8 cartridges (Waters) and the peptide concentration was determined using Pierce colorimetric peptide assay (Thermo Fisher Scientific). The desalted peptides were dried and stored at −20 °C until further use.

### Strong cation exchange chromatography (SCX)

500 μg peptides were loaded onto a PolySULFOETHYL A column (PolyLC INC.) on an Agilent 1260 Infinity II system and separated with a 90 min gradient. All fractions were desalted using C8 StageTips.

### LC-MS/MS analysis

Desalted SCX fractions and high-pH separated enriched phospho peptides were resuspended in 1% ACN, 0.05% TFA and 1 μg peptides were injected into a Thermo Scientific™ Dionex™ UltiMate™ 3000 system connected to a PepMap C-18 trap-column (0.075 mm x 50 mm, 3 μm particle size, 100 Å pore size (ThermoFisher Scientific)) followed by an in-house packed C18 column for reverse phase separation (Poroshell 120 EC-C18, 2.7 μm (Agilent Technologies)) at 300 nL/min flow rate. Peptides were separated using a 180 min gradient and analyzed on an Orbitrap Fusion Lumos mass spectrometer with FAIMS Pro™ device (Thermo Scientific) and Instrument Control Software version 3.4. MS1 and MS2 scans were acquired in the Orbitrap with a mass resolution of 120,000 and 60,000 respectively. MS1 scan range was set to m/z 375 – 1600, standard AGC target, 50 ms maximum injection time and 60 s dynamic exclusion. MS2 scans were set to an AGC target of 1e5, 118 ms injection time, isolation window 1.6 m/z. Only cross-linked precursors at charged states +4 to +8 were subjected to MS2. Peptides were fragmented using stepped collision energy SCE 27 ± 6%. Data were acquired using 2 s per CV with an internal stepping of CVs from -50 to -60 and -75.

### SCOUT

Scout is a user interface-controlled software that accepts many input formats such Orbitrap (.RAW, including files from Orbitrap Astral) or timsTOF (.d) data, and offers a variety of export formats, report of decoys, graphically annotated spectra, statistical analyses, and input tables for xiVIEW^21^ and XlinkCyNET^22^. A full description of the Scout workflow, scores and FDR filtering is provided in Supplementary Notes 1-4.

### Data analysis

RAW files were searched for cross-links with multiple search engines: XlinkX node in Proteome Discoverer v2.5 (Thermo Scientific), MeroX, MaxLynx, MSAnnika, xiSEARCH/xiFDR, and Scout v1.4.1. Search parameters were used as following: MS1 mass tolerance, 10 ppm; MS2 mass tolerance, 20 ppm; maximum number of missed cleavages, 3; minimum peptide length, 6; peptide-mass, 500 – 6,000 Da. Cross-links were detected by searching for the cross-linker modification on lysines. Carbamidomethylation (+57.021 Da) on cysteines was used as a static modification. Oxidation of methionine (+15.995 Da) was set as a variable modification. Data was searched against databases consisting of the proteins that were used to generate the synthetic dataset and differing numbers of random human entries. Wherever applicable, a naïve FDR of 1% (on CSM, XL, and PPI level) was used to filter data (see Supplementary Table 2 for all search settings). Those search parameters are referred to as “standard parameters”. For software tools that did not provide naïve FDRs on all identification levels, naïve FDR-filtered identifications from the next lowest level were aggregated to the higher level without any further post-processing filters and score cutoffs, using the highest reported score of the lower-level identifications as the score for the aggregated higher-level identification.

For the benchmarking of XlinkX, data were post-processed by applying XlinkX score cutoffs. We either applied static score cutoff of 60, as previously recommended^19^, or a dynamic score cutoff as suggested by Thermo (https://assets.thermofisher.com/TFS-Assets/CMD/Reference-Materials/pp-001448-ov-proteome-discoverer-presentation-software-application-pp001448-na-en.pdf), where the cutoffs are selected separately for each dataset based on the highest scoring decoy CSMs.

To enable benchmarking of xiSEARCH, a subset of the dataset had to be analyzed on a computer cluster, using the following parameters suggested by the developers: recalibration of the RAW files; MS1 mass tolerance, 3 ppm; MS2 mass tolerance, 6 ppm; maximum number of missed cleavages, 2. Cross-links were detected by searching for the cross-linker modification on lysines. Carbamidomethylation (+57.021 Da) on cysteines was used as a static modification. Oxidation of methionine (+15.995 Da), deamidation of asparagine (+0.983467), methylation of aspartic and glutamic acids (+14.01565) were set as variable modifications. Search results were FDR-filtered only on PPI-level using xiFDR version 2.1.5.2. The xiFDR boost-function was activated between protein-pairs, remaining settings were left default.

Plot generation and the related data processing were done in the R statistical language.

## Data Availability

The mass spectrometry proteomics data have been deposited to the ProteomeXchange Consortium via the PRIDE partner repository with the dataset identifier PXD042173.

## Code Availability

The Scout software, source code and user documentation are available at https://github.com/diogobor/Scout.

## Acknowledgements

The authors thank Lars Mühlberg, Ying Zhu, Boris Bogdanow, Abigail Lewis and Tara Bartolec for extensive beta-testing and valuable discussion about features to be added. We are grateful for the advice and support from Francis O’Reilly and Anthony Ciancone on how to operate xiSEARCH/xiFDR. F.L. thanks the funding provided by the European Research Council (ERC) Starting Grant (ERC-STG No. 949184) (MR) and by Leibniz-Wettbewerb P70/2018 (FL). FCG is grateful for the support from Fapesp (2014/17264-3) and National Institute of Science and Technology in Bioanalytics (INCTBio). PCC thanks support grants from Fiocruz Inova Produtos Fiocruz PEP, and CNPq.

## Author Contributions

MAC, LUK, PCC, and DBL developed Scout. MR designed and prepared the benchmarking dataset. TC provided recombinant proteins. MR and FCG tested the software and assisted selection and optimization of features. PCC and FL supervised the research and provided funding. MR, FL, MAC, DBL and PCC wrote the manuscript.

## Competing Interests

FL is a shareholder and advisory board member of Absea Biotechnology Ltd., and an advisory board member of VantAI. TC is the co-founder of Absea Biotechnology Ltd. SW is an employee of Absea Biotechnology Ltd. The remaining authors have no competing interests.

